# The genomic basis of heat stress adaptation in indigenous Ethiopian cattle from arid and hot-humid climates

**DOI:** 10.64898/2026.02.09.704906

**Authors:** Endashaw Terefe, Gurja Belay, Abdulfatai Tijjani, Jianlin Han, Bashir Salim, Olivier Hanotte

## Abstract

African zebu cattle (Bos indicus) exhibit remarkable adaptation to extreme thermal conditions, yet the genomic bases of this resilience are not fully elucidated. Ethiopia provides a unique natural setting where closely related zebu populations have divergently adapted to hot-arid (DHETZ) and hot-humid (HHETZ) climates. In this study, we performed whole-genome sequencing of 46 Ethiopian zebu cattle from five populations and compared them with Asian zebu, Sudanese zebu, African taurine, and European taurine breeds. By integrating genome-wide SNP analysis, population genetic structure assessment, and multiple selection scans (*iHS*, *Hp*, *XP-EHH*, and *XP-CLR*), we identified distinct and shared selection signatures between DHETZ and HHETZ cattle. Ethiopian zebu closely clustered with Sudanese zebu but showed clear divergence from Asian zebu and taurine breeds. Although DHETZ and HHETZ cattle exhibited minimal genetic differentiation, reflecting their shared ancestry, each group displayed unique selection signals. DHETZ cattle showed strong selection in genes involved in oxidative stress regulation, protein folding, mitochondrial function, and vascular remodeling (e.g., *SESN2*, *DNAJC8*, *GRPEL2*, *ABLIM3*, and *AFAP1L1*). In contrast, HHETZ cattle displayed signatures in genes associated with immune responses, energy metabolism, and angiogenesis inhibition (e.g., *MYD88*, *PRKACA*, *PRKACB*, and *WIF1*). Several genes, including *VEGFC*, *TNIP3*, and *DMXL2*, were under selection in both groups, suggesting conserved mechanisms of thermotolerance and reproductive adaptation. These findings reveal a dual pattern of genomic adaptation: while core heat-response pathways are shared, population-specific signatures reflect distinct metabolic and vascular strategies for coping with arid versus humid heat stress. This study provides novel insights into the genomic architecture of environmental adaptation in tropical cattle and offers valuable markers for breeding climate-resilient livestock.

## Introduction

Africa harbors one of the most genetically diverse and phenotypically distinct cattle populations, broadly into African zebu (*Bos indicus*), Sanga (taurine-zebu hybrids), Zenga (Sanga-zebu hybrids), and African taurine (*Bos taurus*) breeds [1,2]. Among these, African zebu cattle are the most abundant and widely distributed, particularly across arid and semi-arid regions [3,4]. These cattle exhibit distinct morphological and physiological adaptations to extreme environments, including a prominent hump, large ears, loose skin, and variable coat colors [5,6]. Such features enhance thermoregulation, water conservation, rendering zebu cattle highly suited to tropical and arid climates [7].

Whole-genome studies have revealed substantial genetic diversity within African cattle, reflecting their adaptation to diverse agro-climatic conditions [8,9]. The complex admixture history of African cattle, particularly in the Horn of Africa, has shaped their genomic landscape, supporting resilience to heat stress, disease pressure, and nutritional challenges [8,10,11]. Environmental stressors such as pathogens, high temperatures, and altitude have driven positive selection on functional genomic regions, enhancing survival in harsh conditions [12]. Indeed, heat stress adaptation in tropical cattle involves an integrated suite of morphological, behavioral, physiological, neuroendocrine, and metabolic modifications [13,14]. For example, increased sweating capacity enhances heat dissipation, while lighter coat color, shorter hair, and reduced body size improve thermotolerance [13,15,16].

At the genomic level, candidate genes associated with heat adaptation have been previously identified in African and other cattle populations [17,18]. These previously identified candidate genes linked to oxidative stress regulation (*SOD1*, *GPX7*, *PLCB1*), sweat gland development, hair-coat modification, and energy homeostasis [15,19]. Heat shock proteins (HSPs) such as *HSF5*, *HSPA9,* and *DNAJC18* stabilize protein function under thermal stress, while the endocrine system, through corticotropin-releasing hormone, adrenocorticotropic hormone, and antidiuretic hormone, regulates thermoregulation, water balance, and metabolic adjustments [20,21]. Notably, the *PRLH* gene and its receptor, *PRLR,* have been linked with the slick hair phenotype, which enhances heat dissipation [13,22].

Ethiopia represents a unique ecological setting where zebu cattle are found in close geographic proximity but are adapted to contrasting environments, ranging from hot-arid lowlands to hot-humid highlands [11,12]. Indigenous Ethiopian cattle are predominantly indicine (*B. taurus indicus*), with the exception of the admixed Sheko breed, which carries the highest taurine ancestry among Ethiopian populations [23]. Despite adaptation to divergent environments, Ethiopian zebu exhibit high genetic relatedness, making them an excellent model for identifying genomic regions under selection for thermotolerance [11,24].

In this study, we analyzed whole-genome sequencing (WGS) data from 46 Ethiopian zebu cattle representing five populations inhabiting low-altitude regions (<1500 m above sea level). Based on their ecological adaptation, these populations were grouped into: Dry-Heat Ethiopian Zebu (DHETZ); Afar, Ogaden, and Boran, adapted to hot-arid conditions, and Humid-Heat Ethiopian Zebu (HHETZ); Mursi and Goffa, adapted to hot-humid conditions [11]. We aimed to identify candidate genomic regions under selection for heat stress adaptation in these two contrasting environments and compare them with other indicine populations. Our findings provide novel insights into the genomic basis of thermotolerance in Ethiopian cattle, with direct implications for the genetic improvement of climate-resilient livestock.

## Materials and Methods

### Agro-climatic characteristics of dry-hot and hot-humid agroecologies

#### Dry-hot environment

A dry hot environment is defined by tropical arid conditions characterized by persistently high temperatures, low humidity, and minimal precipitation, typical of arid and semi-arid climates [25]. The Afar, Ogaden, and Borana plateaus, where Afar, Ogaden, and Boran cattle are predominantly raised, exemplify this agroecology. The annual mean temperature in these areas approximates 30°C, with a seasonal variation from 11°C in January to peaks of 45^°^C in June. Precipitation is scarce and highly variable, ranging from 2 mm in January to 56 mm in April.

#### Hot humid environment

Mursi and Goffa cattle are raised under hot-humid conditions. Goffa cattle are primarily located in the Goffa zone near Arbaminch, while Mursi cattle inhabit the South Omo zone. This geo-ecological region typically experiences relatively stable temperatures of 18°C - 21°C throughout the year, although extremes may fall to 10°C or rise as high as 30°C [25]. Rainfall is substantially higher than in dry-hot zones, with monthly precipitation ranging from 48 mm in January to 522 mm in October. Importantly, this hot-humid climate supports the proliferation of tsetse flies, vectors closely associated with trypanosome transmission and prevalence.

### Whole-genome sequencing data of Ethiopian cattle adapted to lowland environments

We analyzed whole-genome sequencing (WGS) data from 46 Ethiopian cattle representing five populations: Afar (n = 9), Boran (n = 10), Goffa (n = 8), Mursi (n = 10), and Ogaden (n = 9). These cattle are from low-altitude regions (<1500 m above sea level) with distinct climatic pressures. Afar, Ogaden, and Boran cattle are adapted to hot-arid environments and collectively designated as the Dry-Heat Ethiopian Zebu (DHETZ) group. In contrast, Mursi and Goffa cattle thrive under- hot humid conditions, forming the Humid-Heat Ethiopian Zebu (HHETZ) group.

To investigate genetic adaptations, we compared Ethiopian WGS data with those from non-Ethiopian cattle populations: 18 Sudanese zebu (SUZ; *n* = 9 Butana and *n* = 9 Kenana), 27 Asian zebu (ASZ; *n* = 10 Gir, *n* = 3 Bhagnari, and *n* = 2 each from Achai, Cholistani, Dhanni, Gabrali, HisarHirayan, Sahiwal, and Tharparkar), 20 European taurine (EUT; *n* = 10 Angus and *n* = 10 Holstein), and 20 African taurine (AFT; *n* = 10 Muturu and *n* = 10 N’Dama). All WGS datasets were retrieved from publicly available repositories, including the NCBI Sequence Read Archive (SRA) (https://www.ncbi.nlm.nih.gov/sra), the China National Gene Bank (CNGB) Nucleotide Sequence Archive (CNSA) (https://db.cngb.org/cnsa/), and EMBRAPA SEG (Accession: 20.18.01.018.00.00) (S1 Table).

### Read mapping and variant calling

Quality control of raw sequence reads was performed using Trimmomatic v0.38 [26], removing short reads (< 35 bp) and low-quality bases (Phred score < 20). Clean paired-end reads were mapped to the *Bos taurus* reference genome ARS-UCD1.2 [27] using the Burrows-Wheeler Alignment (BWA-MEM) algorithm (BWA v.0.7.17) [28]. Subsequent processing included base quality score recalibration (BQSR) and variant calling with the Genome Analysis Toolkit (*GATK v3.8-1-0-gf15c1c3ef*), following best practices recommendations [29]. To enhance variant accuracy, known polymorphisms from the ARS1.2PlusY_BQSR_v3.vcf.gz dataset provided by the 1000 Bull Genomes Project Run 8 (http://www.1000bullgenomes.com/) were used for marking known sites in the bovine genome. Variants of each sample were identified using the GATK *HaplotypeCaller*, and a joint genotyping approach was applied across all samples. Variant quality score recalibration was performed with a 99.9% truth sensitivity filter to minimize false discovery rate and reduce noise from low-quality calls.

### Variant discovery and population genetic structure

SNPs were identified from the WGS data, and the summary statistics were computed using *vcf-stats* of *vcftools v.0.1.15* [30]. Metrics recorded included the total number of biallelic SNPs, autosomal SNPs per sample, heterozygous-to-homozygous (Het/Hom) ratio, genomic inbreeding, and sequencing depth. These parameters were extracted using the *bcftools v.1.8* software [31]. Population genetic structure was assessed with *Admixture v.1.3.0* [32], estimating ancestry proportion under assumed ancestral backgrounds ranging from *K* = 2 to K = 10. The optimal K-value was determined based on the lowest cross-validation error. Ancestry proportions across different K-values were visualized using the R package. To further investigate population differentiation, principal component analysis (PCA) was performed using *Plink v.1.9* [33], comparing Ethiopian cattle groups (DHETZ and HHETZ) with non-Ethiopian populations (ASZ, SUZ, AFT, and EUT).

### Identification of selection signatures

To detect genomic regions under positive selection in Ethiopian cattle, we employed pooled heterozygosity (*Hp*) and integrated haplotype homozygosity (*iHS*). Comparative genome selection scans were performed between the two Ethiopian cattle groups (DHETZ and HHETZ) and Asian zebu (ASZ) using the cross-population extended-haplotype homozygosity (*XP-EHH*) and composite likelihood ratio (*XP-CLR*) tests, applying a 100 kb sliding window with a 50 kb step size. The *HP* analysis was conducted to identify regions of excess heterozygosity within the combined genomes of DHETZ and HHETZ cattle [34]. The *iHS* statistic was used to detect signatures of selection within the Ethiopian zebu population by comparing the integrated extended haplotype homozygosity (EHH) of ancestral alleles versus derived alleles around a core SNP [35], after phasing and imputing the genotypes using the Beagle *v5.1* software[36]. The *XP-EHH* was applied to compare Ethiopian zebu and ASZ cattle, to detect alleles at high frequency to near fixation in Ethiopian cattle relative to the non-Ethiopian [37]. Additionally, XP*-CLR* was computed to identify genomic regions under selection by measuring allele frequency differences between the Ethiopian and ASZ cattle. The method is highly sensitive to recent selection events and detects deviations from neutrality, indicative of both hard and soft selective sweeps [38].

### Functional annotation and characterization of candidate genes

Genome regions identified through selection scan methods were annotated using the Ensembl Biomart genome browser (https://www.ensembl.org/) based on the ARS-UCD1.2 bovine reference genome, to identify candidate genes. Protein-coding genes detected across the four selection scan methods were merged and further analyzed using the Database for Annotation, Visualization, and Integrated Discovery (DAVID) bioinformatics tool (https://david.ncifcrf.gov/) to assess their involvement in biological processes, molecular function, and Kyoto Encyclopedia of Genes and Genomes (KEGG) pathways [39]. Genes with significant functional enrichment (Fisher’s exact test *P* value < 0.05 and fold enrichment > 1.0) were considered functionally relevant for heat stress adaptation.

## Results

### Variant discovery and population structure

Following stringent quality control of whole genome sequencing (WGS) data, we identified 34.2, 33.7, 29.4, 32.2, 16.1, and 13.8 million biallelic autosomal SNPs in HHETZ, DHETZ, SUZ, ASZ, AFT, and EUT cattle, respectively (Table 1; S1 Table). Of these, 93.4%, 91.7%, 93.0%, 87.5%, 93.5%, and 95.1% were known SNPs, mapped to the ARS-UCD1.2 taurine cattle reference genome. The average number of SNPs per sample ranges from 13.8 million in ASZ to 5.7 million in AFT and EUT cattle, with taurine cattle consistently exhibiting fewer SNPs compared to Zebu populations.

**Table 1.**
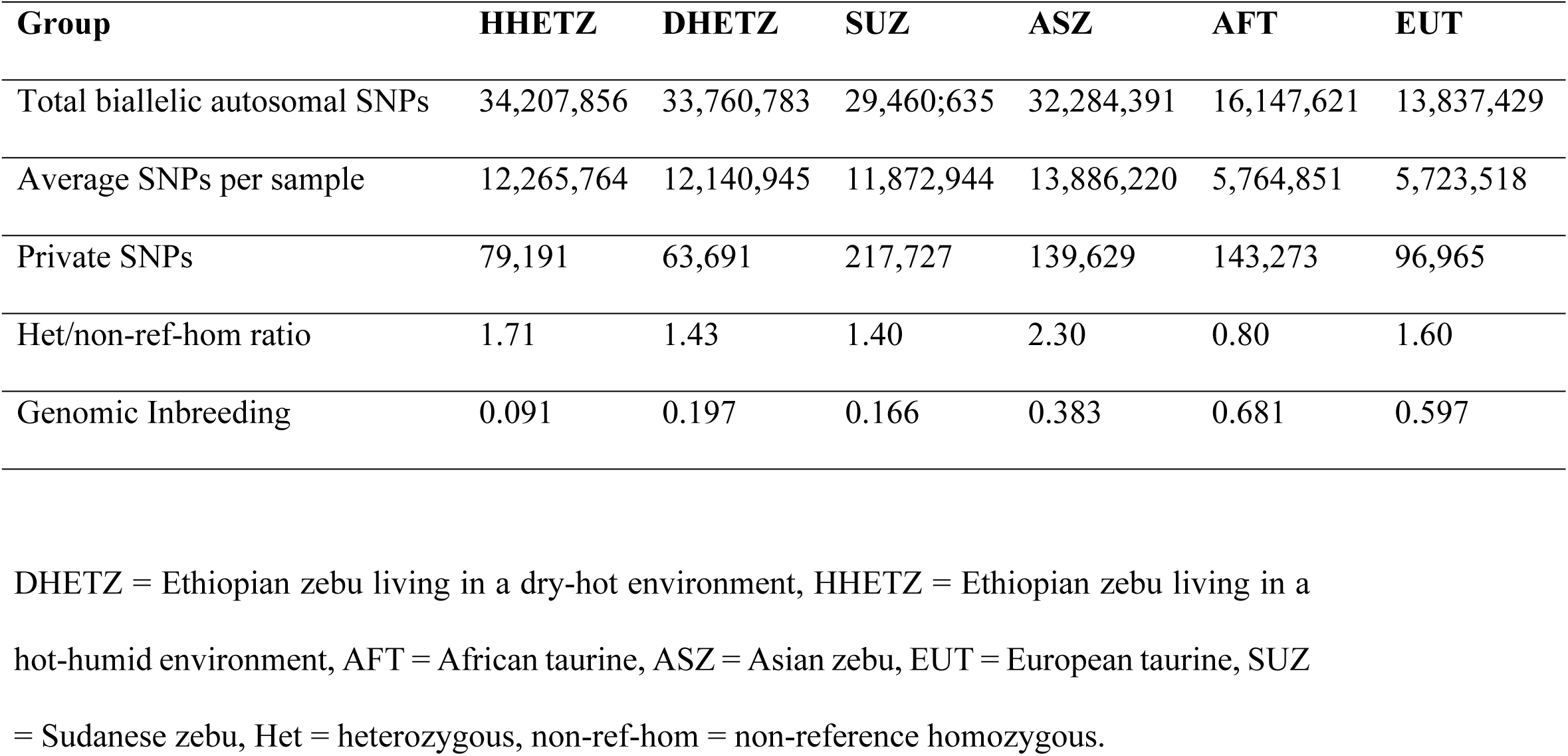
Summary of SNP statistics, including total SNP count, average SNPs per sample, heterozygosity ratio, and genomic inbreeding across the studied cattle populations.

| Group | HHETZ | DHETZ | SUZ | ASZ | AFT | EUT |
| --- | --- | --- | --- | --- | --- | --- |
| Total biallelic autosomal SNPs | 34,207,856 | 33,760,783 | 29,460,635 | 32,284,391 | 16,147,621 | 13,837,429 |
| Average SNPs per sample | 12,265,764 | 12,140,945 | 11,872,944 | 13,886,220 | 5,764,851 | 5,723,518 |
| Private SNPs | 79,191 | 63,691 | 217,727 | 139,629 | 143,273 | 96,965 |
| Het/non-ref-hom ratio | 1.71 | 1.43 | 1.40 | 2.30 | 0.80 | 1.60 |
| Genomic Inbreeding | 0.091 | 0.197 | 0.166 | 0.383 | 0.681 | 0.597 |

DHETZ = Ethiopian zebu living in a dry-hot environment, HHETZ = Ethiopian zebu living in a hot-humid environment, AFT = African taurine, ASZ = Asian zebu, EUT = European taurine, SUZ = Sudanese zebu, Het = heterozygous, non-ref-hom = non-reference homozygous.

Principal component analysis (PCA) explained 61.5% of the total genetic variance across the five cattle groups. PC1 (48.7%) separated the zebu populations (DHETZ, HHETZ, SUZ, and ASZ) from taurine cattle (AFT, EUT), while PC2 (12.8%) distinguished African cattle (DHETZ, HHETZ, SUZ, AFT) from non-African populations (EUT and ASZ) (Fig. 1A). When PC1 and PC3 (7.7%) were analyzed, ASZ and AFT were separated from African zebu and EUT (Fig. 1B). Ethiopian zebu cattle (DHETZ and HHETZ) clustered closely with Sudanese zebu (SUZ), reflecting their shared zebu ancestry and geographical proximity.

Within the Ethiopian cattle, PC1 clearly differentiated DHETZ (Afar, Ogaden, and Boran) from the HHETZ (Mursi and Goffa), consistent with their contrasting agro-ecological backgrounds. However, PC2 did not reveal further sub-structuring among Ethiopian groups (Fig. 1C).

**Fig 1.**
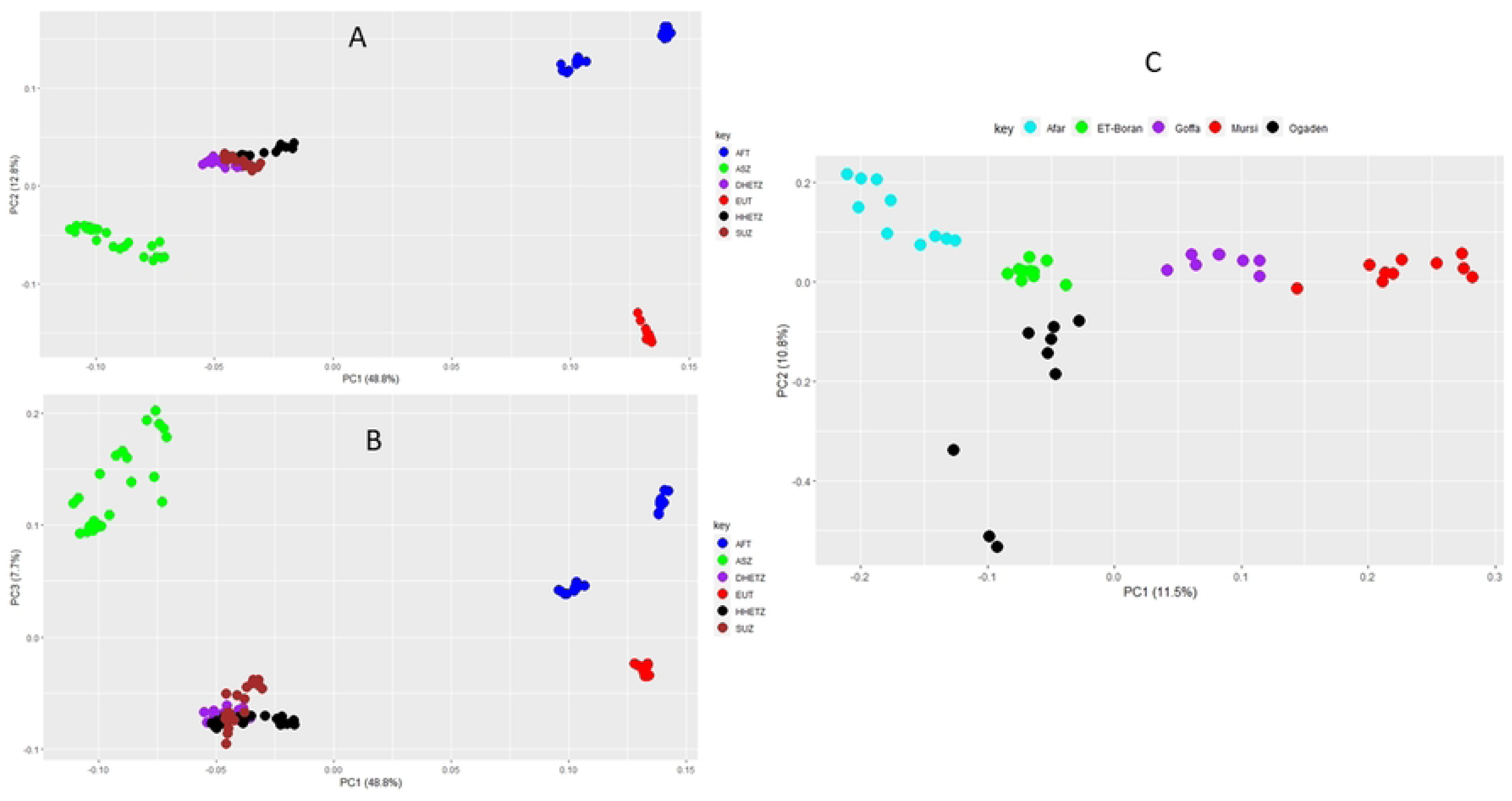
Principal component analysis result. (A) PC1 and PC2 of all populations in the dataset. (B) PC1 and PC3. (C) PC1 and PC2 of Ethiopian cattle populations adapted to a hot-dry environment (Afar, Boran, and Ogaden), and a hot-humid environment (Goffa and Mursi). Population adapted in dry-hot environment (DHETZ), humid hot environment (HHETZ), Sudanese zebu (SUZ), African taurine (AFT), Asian zebu (ASZ), and European taurine (EUT) cattle.

Ethiopian Zebu groups (DHETZ and HHETZ) display comparable SNP counts, despite inhabiting contrasting agro-ecologies (dry-hot vs. hot-humid). In contrast, ASZ cattle display the highest total SNP count and per-sample average, likely reflecting their diverse origins. Both DHETZ and HHETZ harbor fewer private SNPs relative to the reference genome, yet they exhibit higher total SNP counts and heterozygosity-to-homozygosity ratio, consistent with increasing genetic diversity (Table 1; S1 Table).

The heterozygous-to-homozygous ratio varies significantly across populations, with values of 1.71 in HHETZ and 1.43 in DHETZ. These are lower than those in ASZ (2.3) but higher than in AFT cattle (0.8). Ethiopian cattle exhibit lower levels of genomic inbreeding, with HHETZ showing the highest heterozygosity and lowest inbreeding. While SUZ cattle also show elevated heterozygosity, which may reflect recent admixture. DHETZ and SUZ cattle showed comparable heterozygosity and inbreeding.

Population differentiation analyses were performed in *vcftools*/*0.1.15* with 20 samples per group using a 100 kb window and 50 kb step size [30]. It reveals the lowest genetic differentiation between HHETZ and DHETZ (*F_ST_* = 0.0063). Both groups also cluster closely with SUZ cattle, with *F_ST_*= 0.0242 and 0.0231, respectively, reflecting their geographic proximity and shared ancestry. Among Ethiopian groups, DHETZ shows the closest affinity to SUZ, consistent with their common history and adaptation to dry-hot environments (Table 2). A broader comparison showed that African zebu (HHETZ, DHETZ, SUZ) share a more similar genetic structure relative to Asian zebu (ASZ). Across all pairwise comparisons, the highest genetic differentiation was observed between taurine cattle (AFT and EUT) and ASZ (Table 2).

**Table 2.** Population differentiation among the cattle groups in the dataset.

|  | **HHETZ** | **DHETZ** | **SUZ** | **ASZ** | **AFT** |
| --- | --- | --- | --- | --- | --- |
| **DHETZ** | 0.0063 | 0 |  |  |  |
| **SUZ** | 0.0242 | 0.0231 | 0 |  |  |
| **ASZ** | 0.0644 | 0.0572 | 0.0658 | 0 |  |
| **AFT** | 0.2370 | 0.2564 | 0.2500 | 0.3458 | 0 |
| **EUT** | 0.2770 | 0.2933 | 0.2889 | 0.3681 | 0.1946 |

DHETZ = Ethiopian zebu living in a dry-hot environment, HHETZ = Ethiopian zebu living in a hot-humid environment, AFT = African taurine, ASZ = Asian zebu, EUT = European taurine, SUZ = Sudanese zebu.

Genetic admixture analysis was conducted using autosomal biallelic SNPs after removing linked SNPs (LD > 0.5). The optimal number of ancestral clusters is *K* = 4 (Fig. 2A). At this level, Ethiopian zebu (ETZ) includes two indicine-specific ancestries, one African (93.3%) and a minor one shared with Asian zebu (2.8%), and two taurine-specific ancestries, an African taurine (3.4%) and a minor one (0.5%) shared with European taurine. At K = 3, HHETZ displays a relatively higher proportion of taurine ancestry compared to DHETZ (Fig. 2B), whereas DHETZ shares a greater zebu ancestry with ASZ. Additionally, a comparable ASZ component is observed in SUZ cattle compared to DHETZ. At K = 5, the African taurine background, exemplified by the N’Dama, is detected at a higher proportion in HHTZ than in DHETZ cattle.

**Fig 2.**
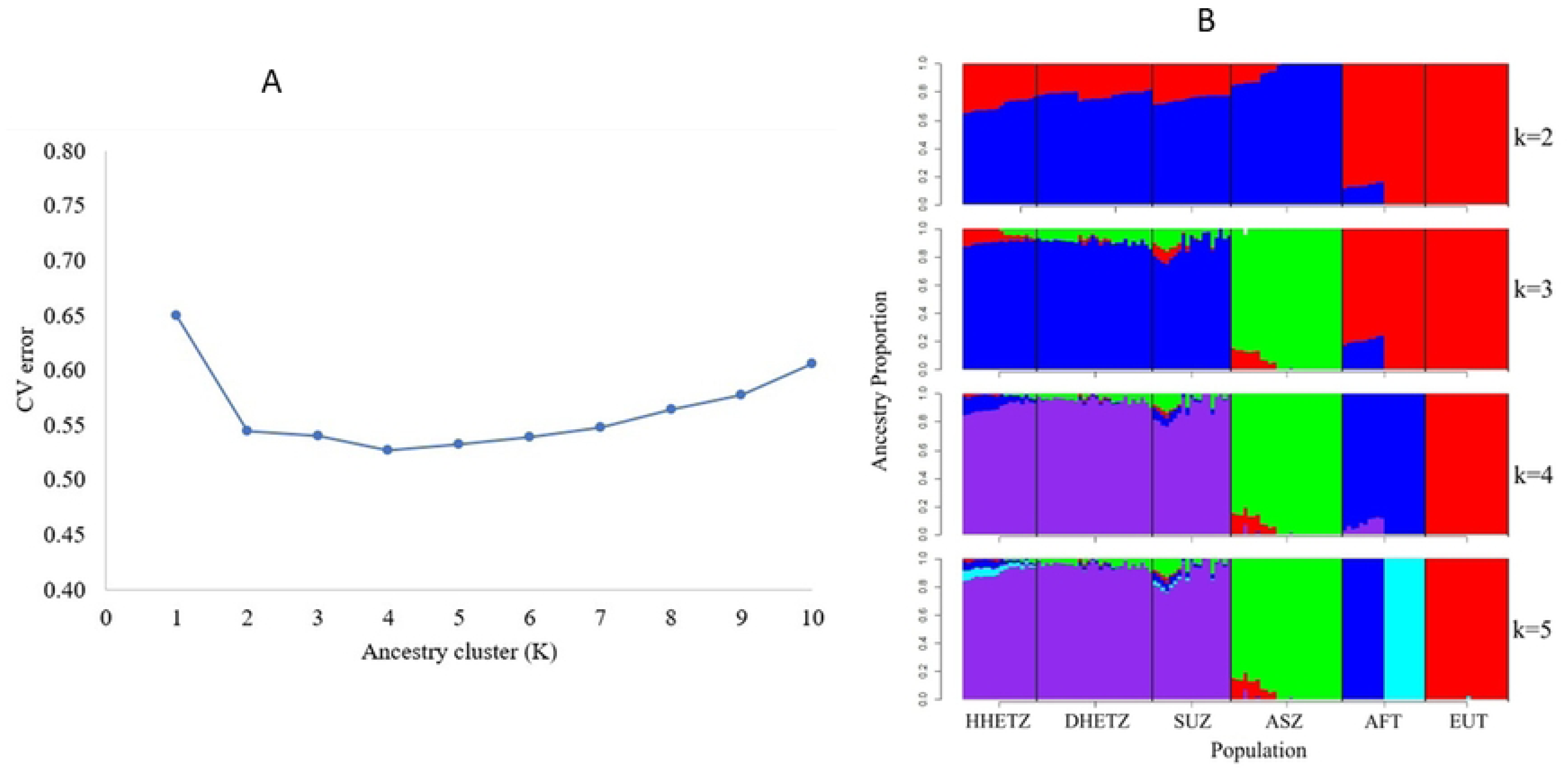
Population genetic structure A) Validation error coefficient all cattle group admixture analysis, B) admixture plot of whole population dataset containing the Ethiopian zebu cattle in a dry hot environment (DHETZ) and hot humid environment (HHETZ), Sudanese zebu (SUZ), African taurine (AFT), Asian zebu (ASZ), and European taurine (EUT) cattle.

### Selection signature within the Ethiopian cattle populations

#### Dry-hot environment (DHETZ)

To identify genomic regions under selection in Ethiopian cattle inhabiting dry-hot environments (DHETZ), we employed integrated haplotype score (*iHS*) and pooled heterozygosity (*Hp*) analyses. A total of 298 non-overlapping genome regions were identified by *iHS*, at a threshold of *P* < 3.96E × 10⁻⁵ (Fig. 3A), encompassing 502 protein-coding genes annotated against the Ensembl ARS-UCD1.2 taurine reference genome (S2 Table). The *HP* scan identifies 113 non-overlapping candidate genome regions under selection at *Z-Hp <-3.3* (Fig. 3B), overlapping with 183 protein-coding genes (S3 Table). Together, *iHS* and *H_P_* detected 33 candidate genes in common, spanning 2.7 Mb across BTA 2, 3, 5, 6, 7, 10, 11, 13, 18, 20, 28, and 29.

**Fig 3.**
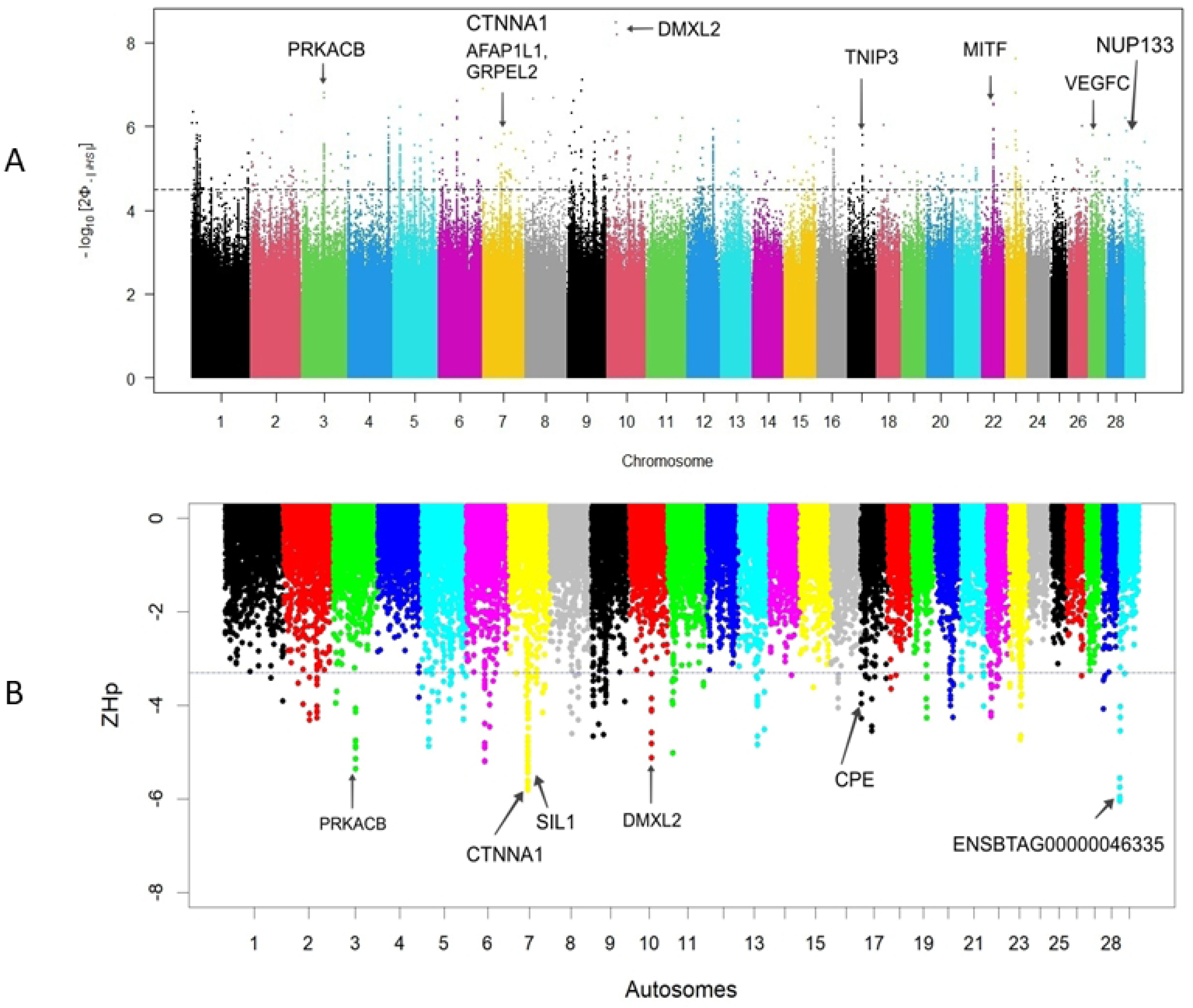
Manhattan plots of genome regions detected by (A) *iHS* and (B) *Hp* in DHETZ cattle. Candidate genes annotated on each plot are genes identified by at least three of the four selection scan methods (*iHS*, *HP*, *XP-EHH*, and *XPCLR*).

#### Hot-humid environment (HHETZ)

Applying the same *iHS* and *Hp* methods to cattle adapted to hot-humid environments (HHETZ) revealed 244 and 138 non-overlapping genome regions, respectively, at a threshold of *iHS* = 5.2 (P value of 6.22 × 10⁻⁶) and ZHp = -3.2 (Figs 4A and B). These regions overlap with 339 (*iHS*) and 183 (*Hp*) candidate genes (S4 and S5 Tables). Twenty-six candidate genes are jointly detected by both methods. These are distributed across BTA 5, 6, 9, 10, 12, 13, 15, and 19. They cover 2.5 Mb of the genome.

**Fig 4.**
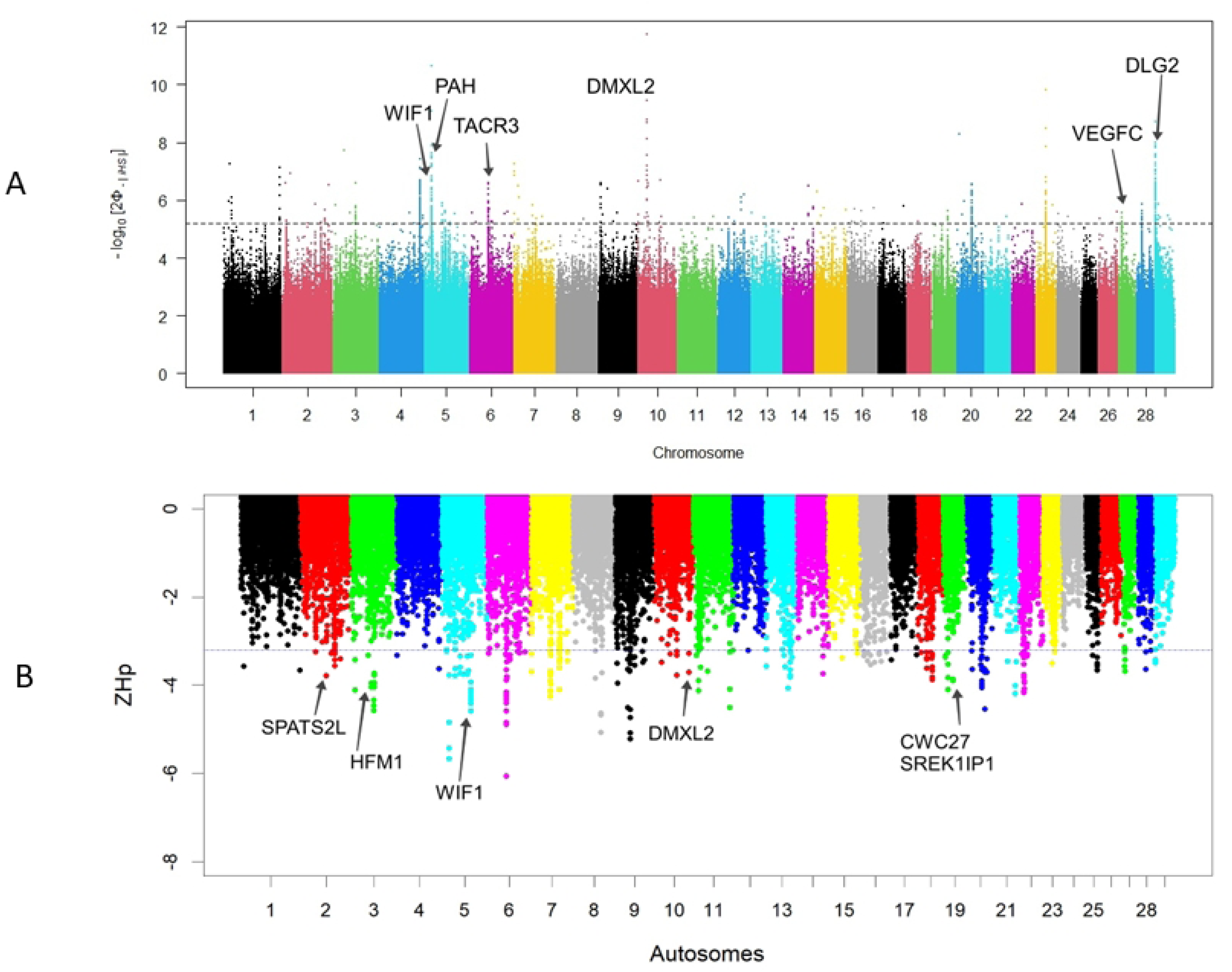
Manhattan plots of candidate genome regions detected by (A) *iHS* and (B) *Hp* selection scan methods in HHETZ cattle. Candidate genes annotated on each plot are genes identified by at least three of the four methods (*iHS*, *Hp*, *XP-EHH*, and *XP-CLR*).

### Comparative genomic signatures of selection among DHETZ, HHETZ, and ASZ populations

#### DHETZ and HHETZ

To identify divergent selection signals between DHETZ and HHETZ populations, we employed cross-population extended haplotype homozygosity (*XP-EHH*) and composite likelihood ratio (*XP-CLR*) analyses. *XP-EHH*, which measures haplotype decay between the populations, detected 163 non-overlapping genomic regions (100 kb window), spanning 27.15 Mb, and harboring 329 protein-coding genes. In parallel, *XP-CLR* identified 227 non-overlapping genomic regions totaling 23.9 Mb and overlapping with 332 protein-coding genes (S6 and S7 Tables). A total of 20 protein-coding genes are shared between *XP-EHH* and *XP-CLR* results. They are localized on BTA 1, 2, 6, 7, 8, 17, 23, and 28 (Table 4). These overlapping signals represent strong candidates for loci underlying adaptive differentiation between cattle inhabiting dry-hot *versus* hot-humid environments.

#### DHETZ and ASZ

Comparative genomic scans between DHETZ and ASZ using *XP-EHH* (*P* < 1.0 × 10⁻⁶) and XP-CLR (score ≥ 90) identified 270 and 207 non-overlapping genome regions, respectively (Figs 5A and 5B). These genome regions were associated with 614 (*XP-EHH*) and 271 (*XP-CLR*) protein-coding genes (S8 and S9 Tables). Across both methods, a total of 56 protein-coding genes were detected. They are located on BTA 2, 3, 4, 5, 6, 7, 10, 11, 12, 16, 17, 22, 23, 27, 28, and 29.

**Fig 5.**
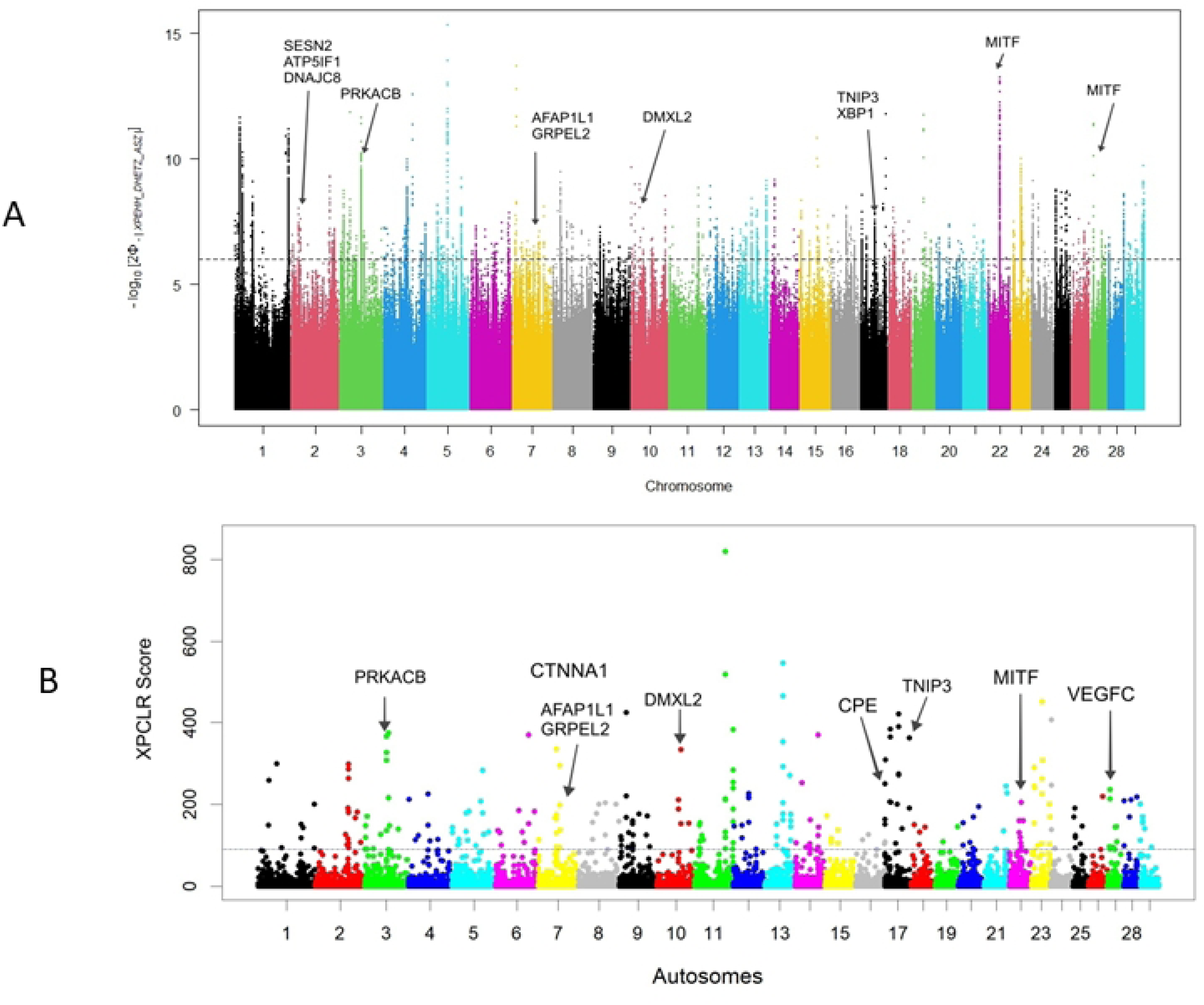
Manhattan plots of genome regions detected between DHETZ and ASZ cattle. (A) *XP-EHH* and (B) *XP-CLR*. Candidate genes annotated on each plot are genes identified by at least three of the four methods (*iHS*, *Hp*, *XP-EHH,* and *XP-CLR*).

#### HHETZ and ASZ

Using the same cross-population tests, 257 and 215 non-overlapping regions were identified between HHETZ and ASZ cattle, within the top 0.5% of 100 kb windows, corresponding to a threshold of *XP-EHH* ≥ 6.0 (*P* < 1.0 × 10⁻⁶) and *XP-CLR* score ≥ 75 (Figs 4C and D). These regions encompassed 644 (*XP-EHH*) and 297 (XP-CLR) protein-coding genes (S10 and S11 Tables). Across both methods, 24 candidate genes are shared. They are distributed across BTA 2, 3, 5, 8, 14, 20, 27, and 29.

**Fig 6.**
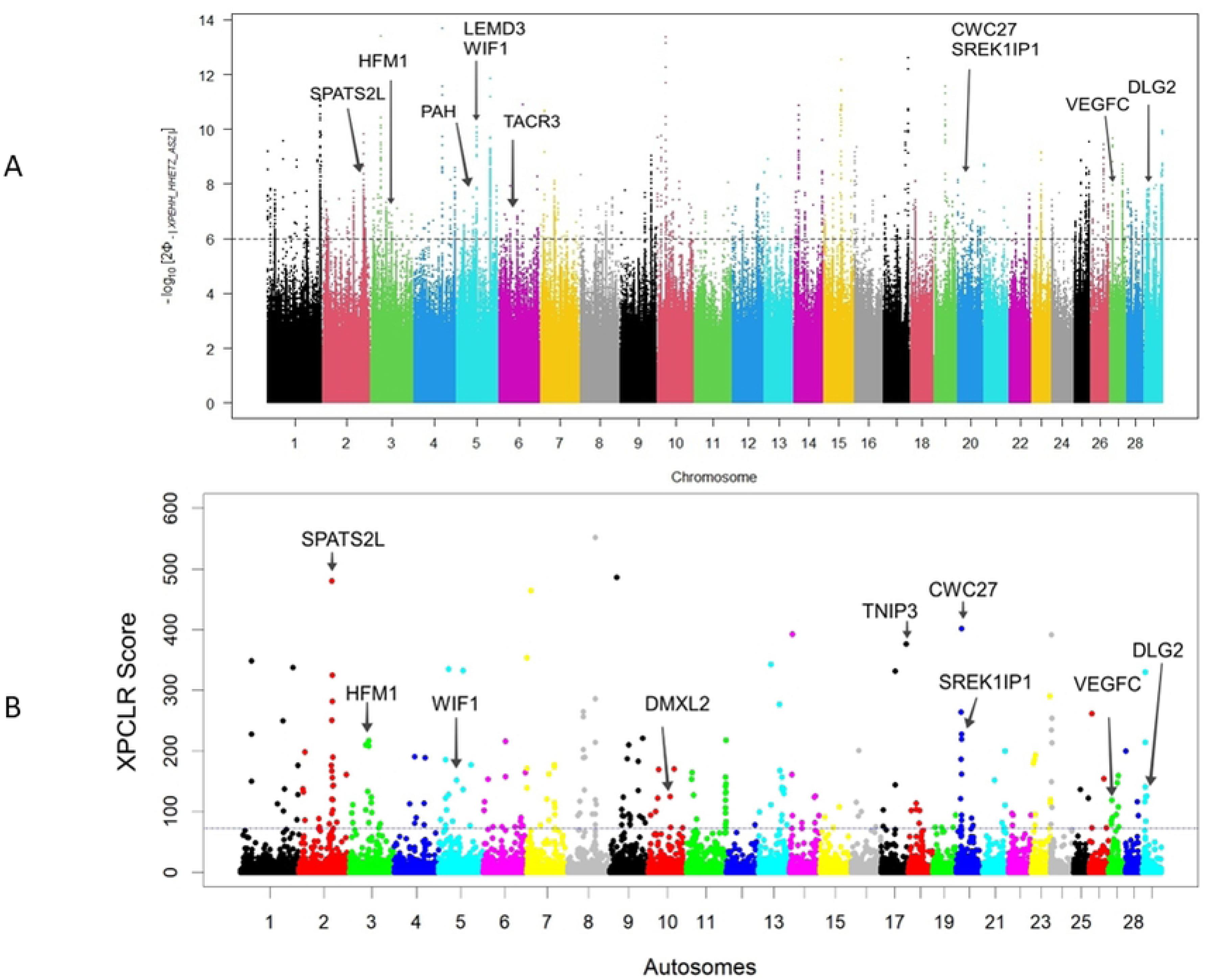
Manhattan plots of candidate genome regions detected between HHETZ and ASZ cattle. (A) *XP-EHH*, and (B) *XP-CLR*. Candidate genes annotated on each plot are genes identified by at least three of the four methods (*iHS*, *Hp*, *XP-EHH*, and *XP-CLR*).

### Overlap selection scans candidate genes and functional annotation

#### DHETZ

Integration of signals from all four selection scans (*iHS*, *Hp*, *XP-EHH*, *XP-CLR*) identified 19 high-confidence candidate regions supported by at least three methods. These regions (2.32 Mb) were distributed across BTA 2, 3, 7, 10, 16, 17, 20, 22, 23, 27, 28, and 29. They include 38 protein-coding genes (Table 2), e.g, *SESN2*, *TP5IF1*, *DNAJC8*, *SAMD13*, *PRKACB*, *TTLL7*, *CTNNA1*, *SIL1*, *ABLIM3*, *AFAP1L1*, *DMXL2*, *GLDN*, *USH2A*, *CPE*, *XBP1*, *TNIP3*, *ADAMTS12*, *MITF*, *VEGFC*, *NUP133*, *ABCB10*, *URB2,* along with several olfactory receptor genes (Table 3). These candidate genes are functionally linked to key traits in adaptation to heat stress, including oxidative stress response, protein folding, heat shock response, melanogenesis, coat pigmentation, reproduction, and angiogenesis.

**Table 3.**
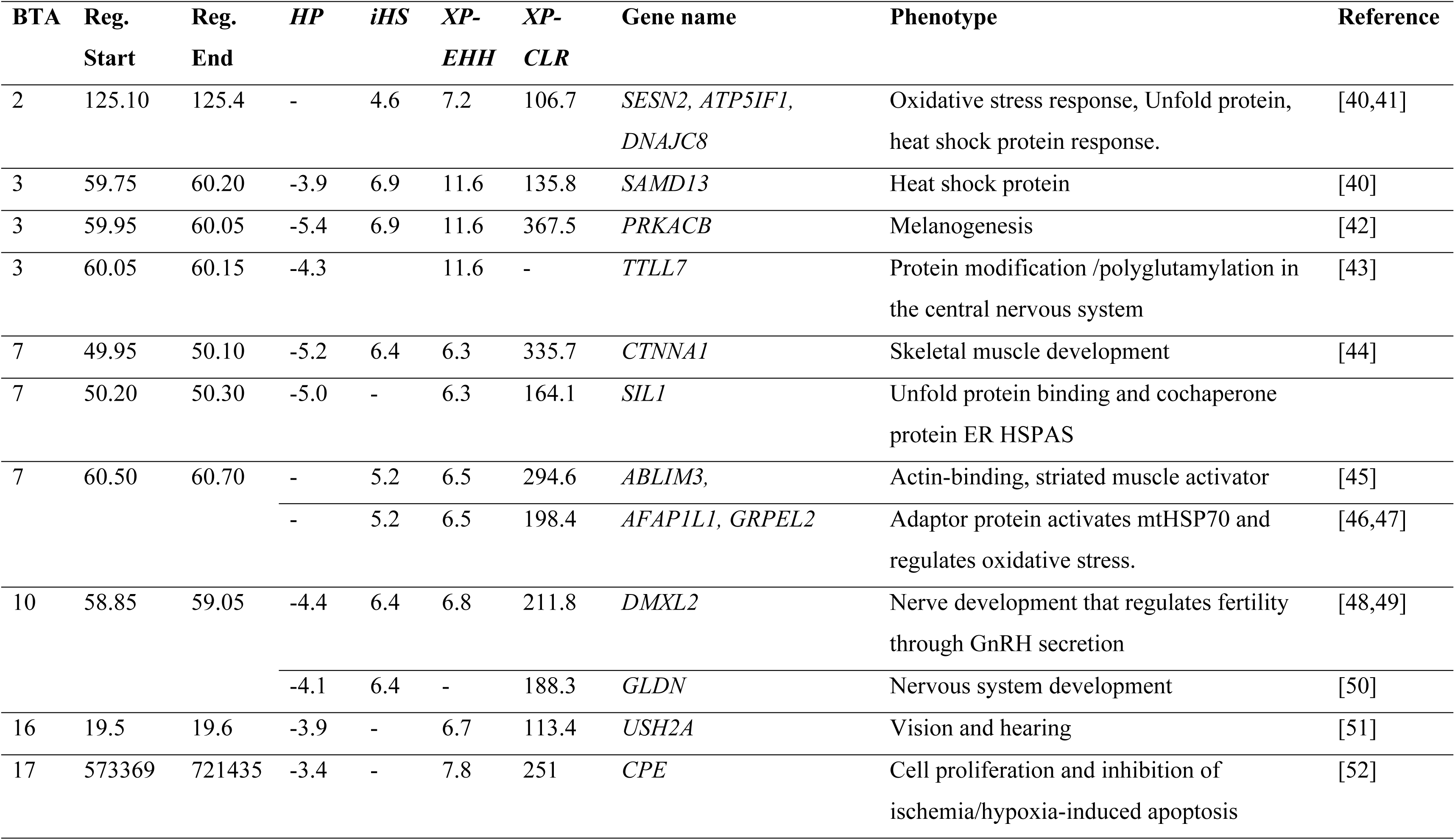
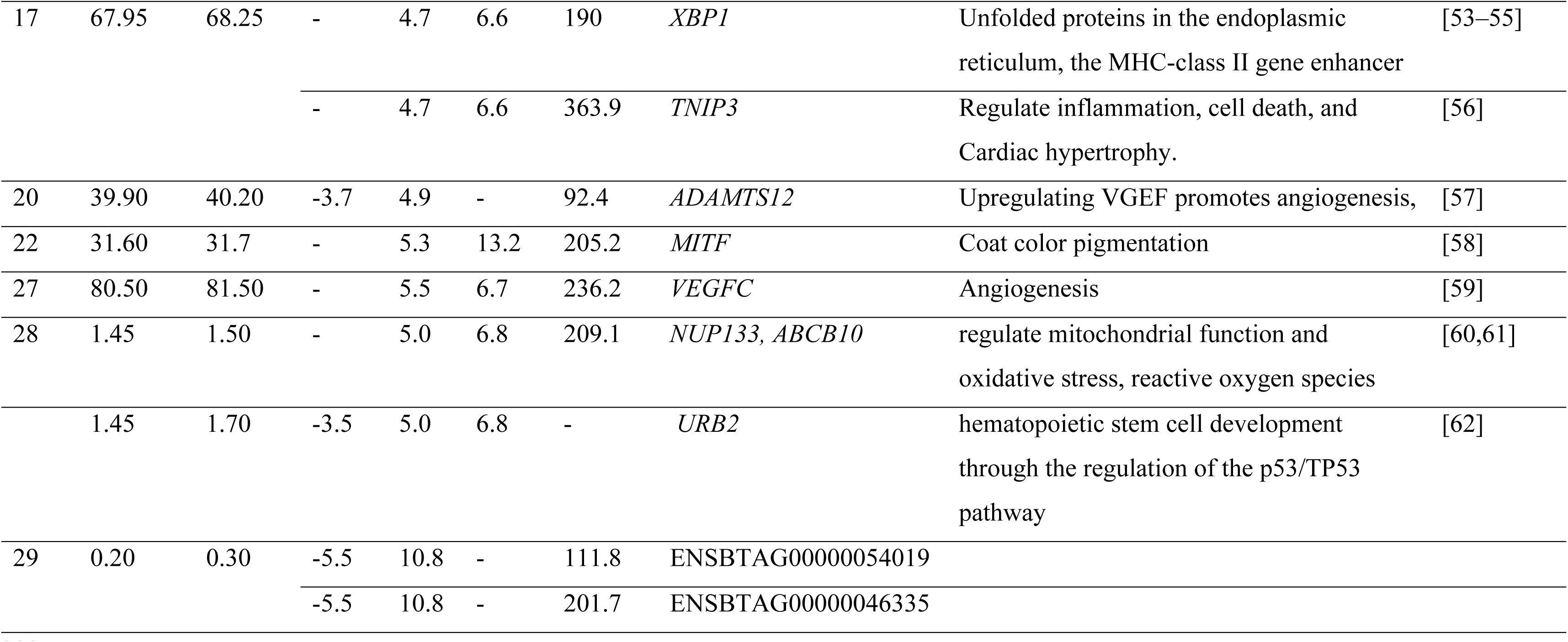
List of candidate regions and associated genes identified in DHETZ cattle by at least three genomic scan methods.

Functional annotation clustering using DAVID revealed enrichment in biological processes such as cellular response to unfolded proteins (GO:0034620), transmembrane receptor protein kinase activity (GO:0071216), response to biotic stimulus (GO:0071216), signal transduction regulation (GO:0009966), and the cellular response to lipopolysaccharide (GO:0071222). Enriched molecular functions included enzyme binding (GO:0019899) and NF-kappa-B binding (GO:0051059). KEGG pathways analysis indicates enrichment in the thyroid hormone signaling pathway (bta04919) and neutrophil extracellular trap formation (bta04613) (S12 Table).

Among the functionally enriched genes, *XBP1*, *ABCB10*, and *TMBIM4* are linked to unfolded protein response and cellular homeostasis during environmental stress. Additional genes such as *XBP1*, *TNIP3*, *PLCG2*, and *IRAK3*, which are involved in lipopolysaccharide response, may contribute to the immune response under stress. Collectively, these genes highlight molecular pathways critical for resilience to heat stress in dry-hot environments.

#### HHETZ

The overlap of candidate genes detected by *HP* and *iHS* within the HHETZ cattle, and by *XP-EHH* and *XP-CLR* between the HHETZ and ASZ, is summarised in Table 4. Among genomic regions under positive selection, 13 candidate regions were identified by at least three of the four methods. These regions, spanning 2.6 Mb across BTA 2, 3, 5, 6, 10, 12, 15, 20, 27, and 29, overlap with 14 protein-coding genes, including *SPATS2L*, *HFM1*, *LEMD3*, *WIF1*, *PAH*, *TACR3*, *DMXL2*, *FRY*, *B3GLCT*, *SOX6*, *CWC27*, *SREK1IP1*, *VEGFC*, and *DLG2* (Table 3). Among these, *WIF1* (WNT inhibitory factor 1) was consistently detected by all four selection scan methods, underscoring its strong selection signal in HHETZ cattle. *WIF1* is located on BTA5, spanning 48.65 - 48.75 Mb. It is functionally associated with anti-angiogenesis, suggesting a potential role in vascular regulation under hot-humid conditions.

**Table 4.** Candidate gene regions and associated genes identified in HHETZ cattle by at least three genomic scan methods.

| BTA | Reg. Start | Reg. End | HP | iHS | XP-EHH | XP-CLR | Gene name | Phenotype | Reference |
| --- | --- | --- | --- | --- | --- | --- | --- | --- | --- |
| 2 | 88.80 | 88.90 | -3.6 | - | 7.1 | 480.2 | *SPATS2L* | Type I interferon, responsible for lupus | [63] |
| 3 | 51.85 | 52.00 | -4.3 | - | 6.9 | 217 | *HFM1* | Gamete formation, Reproduction | [64] |
| 5 | 48.45 | 49.00 | -3.3 | 6.8 | 7.5 | - | *LEMD3* | Regulate cytokine family, like TGF-β, BMPs | [65] |
| 5 | 48.65 | 48.75 | -3.7 | 6.8 | 7.5 | 151.9 | *WIF1* | Anti-angiogenesis | [66] |
| 5 | 66.60 | 66.70 | -3.7 | 7.1 | - | 136.5 | *PAH* | Activate tyrosine to produce melanin | [67,68] |
| 6 | 21.40 | 21.65 | -3.4 | 8.8 | 6.7 | - | *TACR3* | Gonad development and reproduction | [69] |
| 10 | 58.85 | 58.95 | -3.3 | 5.2 | - | 124.9 | *DMXL2* | Fertility | [48,49] |
| 12 | 28.90 | 29.20 | -3.3 | 6.8 | 6.4 | - | *FRY* | Mammary gland development | [70] |
| 12 | 29.60 | 29.75 | -3.9 | 5.5 | 6.8 | - | *B3GLCT* | - | - |
| 15 | 35.70 | 35.85 | -3.4 | 6.5 | - | 73.2 | *SOX6* | Growth and muscle development | [71] |
| 20 | 14.40 | 14.70 | -3.8 | - | 7.2 | 401.9 | *CWC27* | Vision, retinal development | [72] |
|  |  |  | -3.8 | - | 7.2 | 219 | *SREK1IP1* | Activate Vitamin D, related genes | [73] |
| 27 | 7.95 | 8.15 | - | 5.3 | 7.1 | 85.2 | *VEGFC* | Angiogenesis | [59] |
| 29 | 12.05 | 12.20 | - | 5.6 | 7.8 | 125.56 | *DLG2* | Puberty development | [74] |

Gene Ontology (GO) and KEGG pathways enrichment analyses of candidate genes detected in HHETZ populations (S13 Table) highlighted biological processes such as immune response-activating cell surface signaling (GO:0002429), cell surface receptor signaling (GO:0007166), homeostatic process (GO:0042592), and immune response-activating signaling (GO:0002757), Significant molecular functions included glycosyltransferase activity (GO:0016757), and gated channel activity (GO:0022836).

KEGG pathway analysis further revealed enrichment in the thyroid hormone signaling pathway (bta04919), MAPK signaling pathway (bta04010), and African trypanosomiasis (bta05143). Notably, candidate genes *MYD88*, *ENSBTAG00000048268*, ENSBTAG00000050088, and *ENSBTAG00000053508* were grouped within the trypanosomiasis pathway. MYD88 (BTA21) and associated genes on BTA 22 are essential for the innate immune system and immunoglobulin-mediated immune response, potentially enhancing resistance to trypanosome infections in HHETZ cattle.

### Common candidate genes detected between DHETZ and HHETZ populations

A total of nine common genomic regions were identified between DHETZ and HHETZ using the *XP-EHH* and *XP-CLR* (Table 5). These regions contain 18 candidate genes (*MBNL1, DOCK10, RHBDD1, COL4A4, APBB2, NR3C1, RAD50, SMARCB1, DERL3, SLC2A11, MIF, GSTT4, GSTT1, GSTT2, DDT, CABIN1, RAB44*, and *RET*). These loci are distributed across BTA 1, 2, 6, 7, 17, 23, and 28, spanning approximately on1.8 Mb.

**Table 5.**
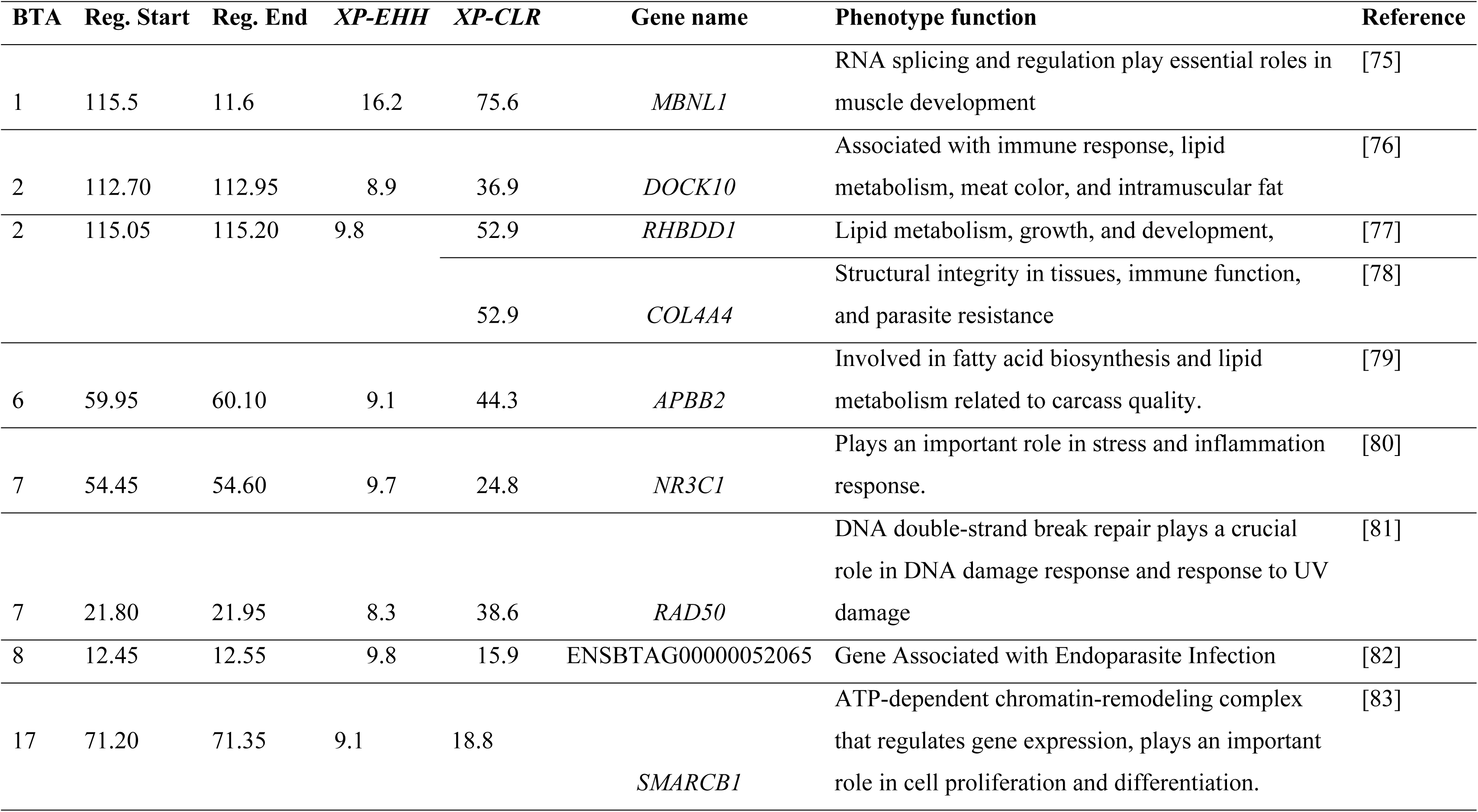
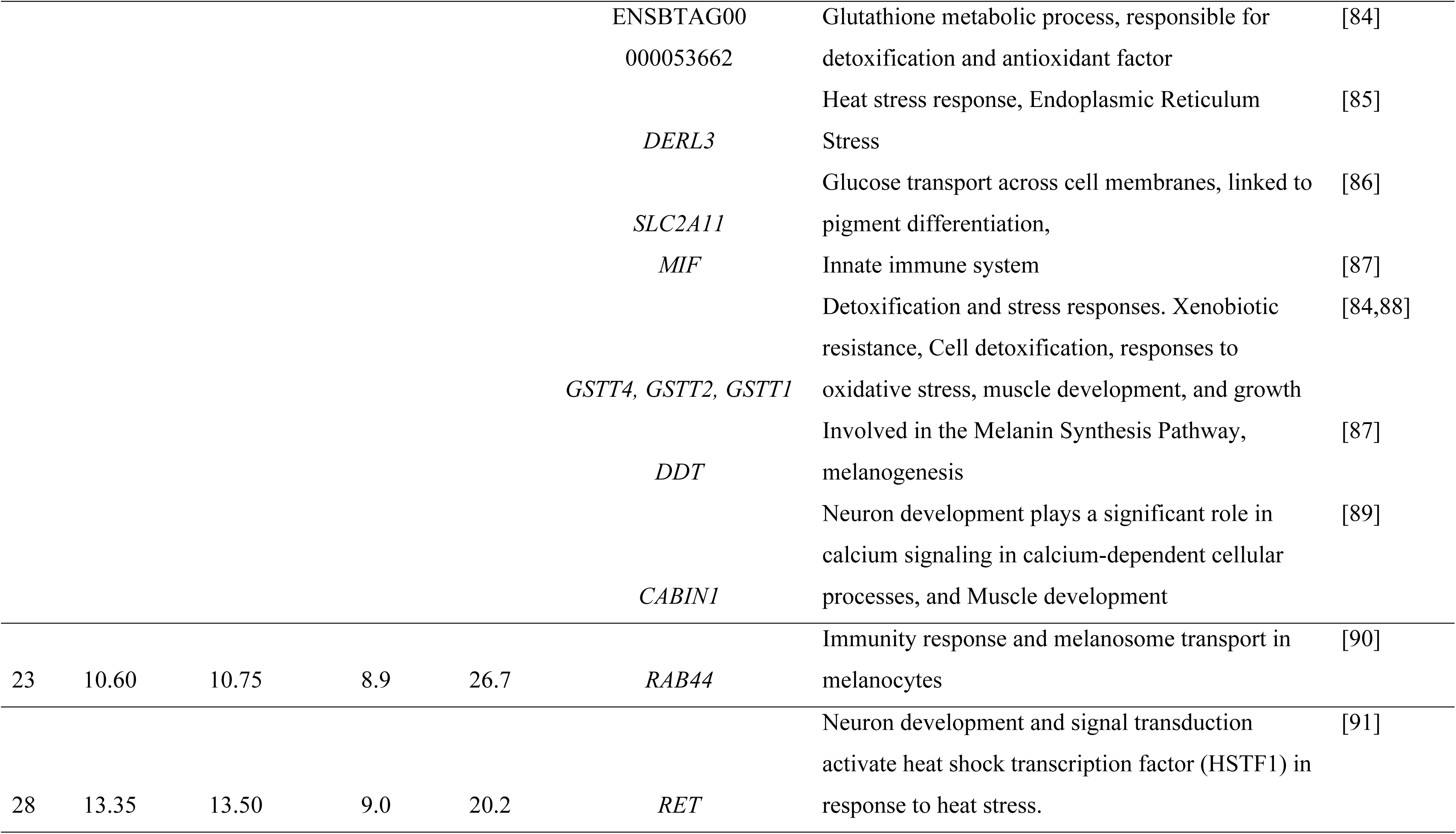
Candidate gene region and associated genes identified in DHETZ and HHETZ cattle.

The functional annotation of these genes reveals significant enrichment in biological processes linked to signal transduction regulation (GO:0009966), reproduction (GO:0022414), and molecular adaptor activity (GO:0060090). KEGG pathways included the PI3K-Akt signaling pathway (bta04151) and melanoma (bta05218), both relevant to stress response and cellular resilience. Several genes (*RAD50*, *NR3C1*, *RHBDD1*, and *KIT*) were implicated in reproductive processes, including meiosis, gametogenesis, and reproductive organ function (S14 Table), underscoring their role in adaptive traits across environmental and developmental contexts.

## Discussion

The ability of African zebu cattle to thrive in extreme thermal environments is driven by genetic adaptations that enhance thermoregulation, oxidative stress tolerance, metabolic efficiency, and immune competence. Deciphering the genetic bases of these traits is critical for sustaining livestock productivity in tropical and arid regions, particularly under the increasing threat of climate change-induced heat stress [25]. In this study, we investigated the genomic architecture of Ethiopian zebu cattle from hot-arid (DHETZ) and hot-humid (HHETZ) environments, comparing their diversity and selection patterns with Asian zebu (ASZ), Sudanese zebu (SUZ), and taurine cattle.

The population structure and admixture analyses reveal that the Ethiopian zebu cattle (DHETZ and HHETZ) share a strong African-specific zebu ancestry and cluster closely with Sudanese zebu. However, they diverge significantly from ASZ and taurine breeds. Principal component and admixture analyses highlight minimal differentiation between DHETZ and HHETZ, suggesting a common ancestral origin. Environment-specific selection pressures or population-specific demographic events might have shaped the small divergence. DHETZ clustered more closely with SUZ, consistent with geography and shared adaptation to arid environments[11]. Overall, our findings emphasize the plasticity of the zebu genome in response to distinct environmental challenges across diverse ecological niches [23,92].

Heat stress can disrupt protein folding, cause cellular damage, impair metabolism, and impact reproductive physiology. To mitigate these effects, molecular chaperones and stress regulators play essential roles in maintaining protein stability and cellular homeostasis [54,93]. In DHETZ cattle, selection scans identified several key candidate genes, including *DNAJC8*, *SAMD13*, *XBP1*, and *ABCB10*, which are critical for protein refolding, mitochondrial protection, and stress-induced endoplasmic reticulum (ER) homeostasis [40,53]. Additionally, candidates include *ABLIM3* and *AFAP1L1* (BTA7:60.6-60.7 Mb), which contribute to muscle fiber integrity, actin cytoskeleton organization, and endothelial proliferation—mechanisms that support vascular remodeling and thermoregulation [45,94]. *ABLIM3*, a member of the actin-binding LIM family of proteins, functions as a cytoskeletal adaptor linking actin filaments to signaling pathways, thereby influencing cell shape, movement, and stress responses [95]. *AFAP1L1*, which encodes an actin filament-associated protein, is involved in muscle development and fat deposition, potentially influencing carcass yield and quality.

The *XBP1* transcription factor is central to the unfolded protein response, regulating genes involved in ER stress recovery, protein folding, and ER-associated degradation. Beyond stress tolerance, *XBP1* also contributes to lipid and glucose metabolism, immune responses, and cellular development [96]. Similarly, *GRPEL2*, a mitochondrial chaperone linked to the *ABLIM3* and AFAP1L1 region, regulates HSP70 activity and ATP-dependent protein folding. By reducing oxidative stress and preserving mitochondrial function, *GRPEL2* enhances resilience to fluctuating temperatures [47,97].

Heat stress also elevates reactive oxygen species (ROS), leading to oxidative damage and metabolic imbalance [98]. The *SESN2* gene, located on BTA2 and selected in DHETZ populations, functions as an antioxidant regulator that maintains redox balance, prevents oxidative stress-induced apoptosis, and safeguards mitochondrial function [99]. These results highlight *SESN2* as a promising biomarker for heat-stress resilience.

In HHETZ cattle, strong selection signals were associated with metabolic adaptation, reproduction, immune defense, and resistance to vector-borne diseases, particularly trypanosomiasis. Metabolic plasticity is a crucial adaptive mechanism for efficient energy utilization under heat stress [5]. Genes such as *PRKACA* and *PRKACB* regulate lipolysis in adipocytes, mobilizing fat reserves to sustain energy production during thermal challenges [100,101].

Immune-related genes under selection included *MYD88* and three uncharacterized loci (*ENSBTAG00000048268*, *ENSBTAG00000050088*, and *ENSBTAG00000053508*), all associated with innate immunity and defense against trypanosome infection. The hot-humid Mursi cattle habitat, savanna grassland with high tsetse fly prevalence [102], likely exerts strong selection pressure on these immune pathways. *MYD88* encodes an adaptor protein activating Toll-like and interleukin-1 receptor signaling, central to immune responses against intracellular pathogens such as *Trypanosoma* spp. [103,104] and *Toxoplasma* spp. [105].

Several candidate genes were detected in both DHETZ and HHETZ, including *DMXL2*, *TNIP3*, and *VEGFC*, reflecting conserved selection for thermotolerance to oxidative stress resilience and metabolic efficiency. For instance, *TNFAIP3* (TNIP3 interacting protein 3) is involved in signal transduction and immunoregulatory, particularly cellular responses to lipopolysaccharide; *VEGFC*, a key regulator of angiogenesis and vascular remodeling [106], enhancing blood circulation and heat dissipation, which is critical for maintaining oxygen delivery during heat stress. In contrast, *WIF1* (WNT inhibitory factor 1), uniquely under selection in HHETZ, acts as an anti-angiogenic factor, inhibiting endothelial cell function through the Wnt signaling pathway [107,108]. This suggests divergent vascular strategies in cattle adapted to arid versus humid environments.

Heat stress also compromises reproductive success, necessitating robust genetic adaptations. Here, *DMXL2*, identified in both groups, regulates gonadotropin-releasing hormone (GnRH) and is essential for ovarian follicle development and reproductive efficiency [49]. Similarly, *HFM1,* detected in HHETZ cattle, is required for spermatogenesis and gametogenesis [64,109]. These findings highlight the genetic foundation of reproductive resilience, ensuring fertility under thermal stress.

Finally, *RET*, identified under selection in HHETZ, is clustered in MAPK signaling and protein kinase pathways, regulating the induction of heat shock protein (*HSP70* family: *HSPA1A*, *HSPA1B*, and *HSPA1L*). These stress-induced molecular chaperones maintain mitochondrial integrity during heat challenges [91].

## Conclusions

This study provides a comprehensive genomic assessment of Ethiopian zebu cattle, uncovering genetic signatures linked to thermoregulation, oxidative stress resistance, metabolic adaptation, and reproductive resilience. Although DHETZ and HHETZ cattle share core adaptive mechanisms, environment-specific selection pressures have driven functional divergence, particularly in metabolic and vascular regulation. These insights advance our understanding of the genetic basis of climate resilience in livestock, with direct implications for sustainable conservation and genomic-informed breeding programs. Future research integrating functional genomics and transcriptomics will be essential to unravel the complex gene-environment interactions shaping thermally adapted livestock populations.

## Supporting information

S1 Table. List of Breed groups, Sequence sources (Bioproject and Biosamples), and SNP analysis in the all cattle dataset. (xlsx)

S2 Table. Candidate genes detected by the integrated haplotype score (*iHS*) selection scan method in DHETZ cattle. (xlsx)

S3 Table. Candidate genes detected by the pooled heterozygosity (*Hp*) selection scan method in DHETZ cattle. (xlsx)

S4 Table. Candidate genes detected by the integrated haplotype score (*iHS*) selection scan method in HHETZ cattle. (xlsx)

S5 Table. Candidate genes detected by the pooled heterozygosity (*Hp*) selection scan method in HHETZ cattle. (xlsx)

S6 Table. Candidate genes detected by the cross-population extended haplotype homozygosity (*XP-EHH*) selection scan method comparing DHETZ and HHETZ cattle. (xlsx)

S7 Table. Candidate genes detected by the cross-population composite likelihood ratio (*XP-CLR*) selection scan method comparing DHETZ and HHETZ cattle. (xlsx)

S8 Table. Candidate gene regions and annotated genes detected by the *XP-EHH* selection scan method in DHETZ and ASZ cattle. (xlsx)

S9 Table. Candidate gene regions and annotated genes detected by the *XP-CLR* selection scan method in DHETZ and ASZ cattle. (xlsx)

S10 Table. Candidate genes detected by the *XP-EHH* selection scan method in HHETZ and ASZ cattle. (xlsx)

S11 Table. Candidate genes detected by the *XP-CLR* selection scan method in HHETZ and ASZ cattle. (xlsx)

S12 Table. The DAVID functional annotation of candidate genes identified by the *Hp*, *iHS*, *XP-EHH*, and *XP-CLR* selection scan methods in DHETZ. (xlsx)

S13 Table. The DAVID functional annotation of candidate genes identified by the *Hp*, *iHS*, *XP-EHH*, and *XP-CLR* selection scan methods in HHETZ. (xlsx)

S14 Table. The DAVID functional annotation of candidate genes identified by the *XP-EHH* and *XP-CLR* selection scan methods comparing DHETZ and HHETZ cattle. (xlsx)

## Acknowledgment

The authors would like to acknowledge the support of the International Livestock Research Institute (ILRI) of the Livestock Genetic Group (LiveGene) and the Center for Tropical Livestock Genetics and Health (CTLGH) African cattle Genetic Research project for supporting the study and funding a PhD research fellowship. We also extend our appreciation to King Faisal University and the CAAS-ILRI Joint Laboratory on Livestock and Forage Genetic Resources in Beijing, China, for their collaborative contributions. This study was part of Endashaw Terefe’s PhD research at Arsi University, supported by ILRI.

